# Unraveling cooperative and competitive interactions within protein triplets in the human interactome

**DOI:** 10.1101/2025.06.10.658265

**Authors:** Aimilia-Christina Vagiona, Pablo Mier, Miguel A. Andrade-Navarro

**Affiliations:** Insitute of Organismic and Molecular Evolution, Faculty of Biology, Johannes Gutenberg University, Biozentrum I, Hans-Dieter-Hüsch-Weg 15, 55128 Mainz, Germany; Andalusian Center for Developmental Biology (CABD, UPO-CSIC-JA), Faculty of Experimental Sciences (Genetics Area), Universidad Pablo de Olavide, 41013 Seville, Spain

## Abstract

Knowledge about protein-protein interactions (PPIs) is crucial for our understanding of cellular functions. An important contribution to these comes from experimental high-throughput techniques such as yeast two-hybrid (Y2H), which provide evidence for direct, binary PPIs. Consequently, although protein function often emerges from interactions within multi-protein assemblies, most PPI networks focus on binary relationships. Higher-order motifs, such as protein triplets, can reflect cooperative or competitive binding events crucial for cellular function. However, distinguishing these interaction types systematically from a network of binary PPIs remains challenging. To address this issue, we present a computational framework to predict cooperative and competitive interactions among protein triplets in the human Protein Interaction Network (hPIN). By embedding the hPIN into two-dimensional hyperbolic space using the LaBNE+HM algorithm, we extracted latent geometric coordinates for each protein. Combined with topological features and biological annotations, these were used to train a Random Forest classifier on structurally validated triplets from Interactome3D. Our model achieved strong performance (AUC = 0.88), identifying angular and hyperbolic distances as two of the most predictive features. Cooperative triplets were characterized by longer common proteins and a lower frequency of paralogous partners, indicating distinct structural and evolutionary features compared to competitive cases. Structural evaluation using AlphaFold 3 confirmed that cooperative triplets exhibit distinct binding sites, whereas competitive ones share overlapping interfaces. Hyperbolic embeddings, in combination with machine learning, offer a powerful approach to characterize higher-order interaction motifs in protein networks. Our findings provide mechanistic insights into the organization of protein complexes and a predictive framework to support future studies in network biology.

## Introduction

Proteins are the essential units of life, regulating crucial biological processes ranging from molecular transport to signal transduction [1]. These functions depend on a highly coordinated network of protein–protein interactions (PPIs), collectively referred to as the interactome, which controls cellular organization and functionality [2,3]. Understanding these interactions is key to unraveling biological mechanisms and designing therapeutic strategies for various diseases [4,5].

Protein-protein interaction networks (PPINs) are primary constructed from binary interactions, which represent direct physical contact between two proteins [6]. Most large-scale interactome maps rely on pairwise interaction data, as these are the most accessible and well-characterized through traditional experimental approaches [7]. High-throughput techniques such as yeast two-hybrid (Y2H) assays and affinity purification coupled with mass spectrometry (AP-MS) have been essential in mapping the human interactome [8,9]. However, many biological processes extend beyond pairwise PPIs, as proteins often function in multi-protein assemblies rather than simple one-to-one interactions, in order to carry out essential tasks [10,11]. Among these multi- protein complexes, protein triplets represent a crucial class of higher-order interactions. Triplet interactions provide a framework for understanding cooperative and competitive dynamics, which influence the structural and functional stability of protein complexes [12,13].

Cooperative interactions occur when multiple proteins work together synergistically to enhance stability or function, such as in multiprotein enzyme complexes or transcription factor binding events [14,15]. In contrast, competitive interactions arise when two proteins compete for the same binding partner, modulating signaling pathways or enzymatic activity based on cellular conditions [16]. Accordingly, proteins interacting with many partners can have either multiple interfaces or just one interface. Some of these interfaces are shared by different partners, resulting in mutually exclusive bindings; other interfaces are used by only one partner such that interactions with different partners can occur simultaneously. Thus, to judge whether two partners can interact with the common protein simultaneously, the key is to know whether they share an interaction interface [17].

Despite their importance, higher-order interactions remain difficult to study using traditional experimental methods, which are primarily designed to detect pairwise interactions. This limitation highlights the need for computational approaches that can infer triplet interactions and predict whether they are cooperative or competitive. By integrating network properties and advanced mathematical models, such as hyperbolic embeddings, researchers can gain a more comprehensive understanding of how protein complexes form and function in the cellular environment [18].

Several studies have demonstrated that complex networks, including the human protein– protein interaction network (hPIN) can be effectively modeled using hyperbolic geometry [19– 21]. The Popularity-Similarity (PS) model provides a geometric framework in which network nodes are positioned within a two-dimensional hyperbolic space (H^2^), represented as a disk. In this model, the radial coordinate of a node captures its popularity and evolutionary age, with older, highly connected proteins positioned closer to the center, while newly emerging proteins occupy the periphery. The angular coordinate encodes functional similarity, grouping proteins involved in shared biological processes or pathways. Alanis-Lobato et al. demonstrated that embedding the hPIN in hyperbolic space provides biologically meaningful insights, with radial positioning reflecting protein conservation and seniority, and angular positioning capturing functional and spatial organization within the cell [22]. Such mapping approaches can contribute to a deeper understanding of complex human disorders [23–25].

Motivated by the importance of higher-order protein interactions, we developed a computational framework to distinguish cooperative from competitive triplets in the human protein interaction network (hPIN). Starting from a high-confidence PPI network embedded in hyperbolic space, we identified open triangles, triplets of proteins where two interact with a shared partner but not with each other, and annotated them using structural data. Using this annotated dataset, we trained and evaluated a Random Forest algorithm to classify protein triplets as either cooperative or competitive based on topological, geometric and biological features. To assess the predictive power of our approach, we performed structural validation using AlphaFold 3, confirming distinct spatial configurations between predicted cooperative and competitive triplets. By exploring multi-protein interactions, we provide a deeper insight into how molecular complexes are organized and operate within biological systems.

## Materials and methods

### Human protein interaction network construction

The human protein-protein interaction network (hPIN) used in this study was built using version 2.3 of the Human Integrated Protein–Protein Interaction rEference (HIPPIE) database [26,27]. HIPPIE integrates experimentally supported physical interactions from major expert-curated resources and assigns a confidence score to each interaction based on evidence type, reproducibility, and publication quality. In this study, the hPIN was constructed using interactions with confidence score ≥ 0.71 because the majority of the edges are supported by more than one experiment [18,24]. Self-interactions were removed, and only proteins within the largest connected component (LCC) were kept. The final network comprises 15,319 proteins and 185,791 high-confidence interactions (Supplementary Tables S1 and S2).

### Mapping the hPIN into the hyperbolic space

We embedded the hPIN in the two-dimensional hyperbolic plane using the R package “NetHypGeom”, which implements the LaBNE + HM algorithm [28]. The algorithm combines manifold learning and maximum likelihood estimation to uncover the hidden geometry of complex networks [29,30]. LaBNE + HM expects a connected network as input, typically the LCC. The other components cannot be mapped due to the lack of adjacency information relative to the LCC. The PS model has a geometrical interpretation in hyperbolic space (H^2^) where nodes that join the system connect with the existing ones that are hyperbolically closest to them [20,31]. The network was embedded in H^2^ to infer the hyperbolic coordinates of each protein, with parameters γ = 2.97, T = 0.83, and w = 2π. The 15,319 nodes of the hPIN lie within a hyperbolic disc where the radial coordinate of a node, r_*i*_, represents the popularity dimension with nodes that joined the system first being close to the disc’s center. The angular coordinate, θ_*i*_, represents the similarity dimension [22].

### Identification of cooperative interactions and structural annotations

We then identified all the possible open triangles within the hPIN, resulting in approximately 17 million unique triplets. An open triangle is defined as a triplet of proteins in which a central node (common interactor) interacts with two others (V1 and V2), which do not interact. We focused exclusively on open triangles because in closed triangles (where all three proteins interact), no central or common node can be uniquely defined as all nodes are equivalent. This simplified structure allowed a more effective way of modeling cooperative and competitive interaction types.

To annotate structural support for these triplets, we used PPI data from the Interactome3D database, which includes residue-level annotations of experimentally resolved structures deposited in the Protein Data Bank (PDB) [32]. Model-based predictions and self-interactions were excluded from the analysis. Protein complexes were grouped by PDB ID, and only those involving at least three proteins were considered. Within each complex, all possible protein triplets were generated, and the number of observed pairwise interactions per triplet was counted. Triplets in which exactly two of the three possible pairwise interactions were present were classified as cooperative interactions in protein triplet complexes. We then integrated the Interactome3D-derived cooperative triplets with the full set of hPIN open triangles to determine the final set of positive cases used for model training and evaluation. To avoid overrepresentation of proteins, present in many complexes or in complexes with many subunits, only one validated triplet per common interactor was retained, by keeping the first occurrence based on dataset order (Supplementary Table S3). All remaining open triangles in the hPIN were considered non-cooperative (or competitive).

### Feature extraction and randomization

Each protein triplet in the dataset was annotated with a total of 42 features, capturing both network topology and biological context at the node and edge levels. For every protein within a triplet (common interactor, V1, and V2) we extracted 11 features, resulting in 33 features per triplet. These included the two hyperbolic coordinates (r and theta) and four network centrality measures: Degree Centrality (DC) [33], Closeness Centrality (CC) [34], Betweenness Centrality (BC) [35], and Eigenvector Centrality (EC) [36]. These centralities quantify different aspects of a node’s importance within the network, such as how connected it is (DC), how close it is to all other nodes (CC), how often it lies on shortest paths (BC), and how influential its neighbors are (EC). To incorporate biological relevance, we further assigned each protein with five additional features: the presence of Intrinsically Disordered Regions (IDRs), obtained from the DisProt database v9.7 [37], and the subcellular localization (nucleus, cytoplasm, endomembrane, and multi-localized) retrieved from the Human Protein Atlas database v24.0 [38].

In addition, three more features were computed for the each of the edges between the possible protein pairs in the triplet (common–V1, common–V2, V1–V2): hyperbolic distance (hd), radial difference (rd), and angular difference (thetad), resulting in 9 edge-level features per triplet. To avoid positional bias during model training, the order of V1 and V2 was swapped in a random selection of 50% of the triplets. This ensured that the model learned from meaningful topological and biological properties rather than any artificial ordering pattern. The complete list of features and their descriptions is provided in Supplementary Tables S1 and S2. The complete dataset that was used for the modeling is provided in Supplementary Table S4.

### Model development and evaluation

Model development was performed using the caret package in R [39]. The primary classification algorithm applied was Random Forest (RF) [40], selected for its robustness and effectiveness in handling high-dimensional biological data. To benchmark performance, we additionally trained and evaluated four other machine learning algorithms: Support Vector Machine (SVM) [41], Logistic Regression (LogReg) [42], Decision Tree (DT) [43], and k-Nearest Neighbors (kNN) [44]. The dataset was split into a training set (70%) and a test set (30%), and class imbalance was addressed by applying random undersampling of the majority class during training. Model training was conducted using 5-fold cross-validation repeated 10 times to ensure robust hyperparameter optimization. For the Random Forest model, parameters such as the number of variables randomly sampled at each split (n=22) and the number of decision trees (n=500) were tuned based on cross-validation performance. Model evaluation was carried out on the independent test set using multiple performance metrics, such as Receiver Operating Characteristic (ROC) curves [45] and the Area Under the Curve (AUC) [46] to assess the model’s ability to distinguish between cooperative and competitive interactions. In addition, we calculated accuracy, precision, recall (sensitivity), and F1-score to comprehensively evaluate classification performance, particularly in the context of class imbalance [47]. Finally, we computed feature importance scores using the Random Forest model to identify the most informative topological and biological features contributing to the classification task.

### Homology and length annotations for biological validation of predicted cooperative and competitive protein triplet complexes

Following the model application to the complete dataset of open triangles in the hPIN, we annotated the predicted interactions with additional biological features to support downstream analysis. Specifically, we integrated information on paralogy between V1 and V2 proteins and protein sequence lengths. Paralogy between V1 and V2 was assessed using curated human data (GRCh38.p14) from the Ensembl database release 113 [48]. For each protein in the triplet, sequence lengths were obtained from the UniProtKB database [49]. These values were used to calculate the individual lengths of the common interactor and the average length of the V1 and V2 proteins.

### Structural validation of predicted cooperative and competitive protein triplet complexes using AlphaFold 3

Following the prediction of protein triplet complexes as either cooperative or competitive, we proceeded with structural modeling to evaluate whether the predicted interaction types were supported by structural evidence. For each protein triplet, the individual protein sequences were retrieved from the UniProtKB database [49] and used as input for AlphaFold 3 [50]. Instead of modeling the entire triplet as a single complex, we performed pairwise structure prediction by separately analyzing the common–V1 and common–V2 protein pairs. Specifically, we examined whether V1 and V2 interacted with the same or overlapping regions on the common interactor protein. Overlap in binding sites was interpreted as evidence of potential competitive binding, while distinct binding interfaces were indicative of potential cooperative interactions. We used PyMOL to visualize the predicted structures, which allowed us to manually examine where the proteins were binding and how they were positioned in 3D space [51].

## Results and Discussion

### Construction of the hPIN into the hyperbolic space and structural annotations of cooperative and competitive relationships

To investigate cooperative and competitive interactions between proteins in triplets, where one protein interacts with another two proteins, we first constructed a high-confidence human protein–protein interaction network (hPIN) using experimentally supported data (Figure 1a). For this, we retrieved all human PPIs from the HIPPIE database and filtered interactions with a confidence score ≥ 0.71, as this threshold ensures that the majority of the interactions are validated through multiple independent sources. The resulting network comprised 15,319 proteins and 187,791 interactions. To uncover the latent geometry underlying the hPIN, we embedded the hPIN into the two-dimensional hyperbolic plane (H^2^) using the LaBNE+HM algorithm, which integrates manifold learning with maximum likelihood estimation. Each protein was assigned with a set of hyperbolic coordinates: a radial coordinate (r) representing its topological centrality where a shorter distance to the center corresponds to nodes with higher connectivity, and an angular coordinate (theta) indicating its similarity to other nodes based on interacting partners [18,22,25]. This hyperbolic embedding allowed us to extract geometric and topological features essential for the classification of cooperative versus competitive interactions within protein triplets.

**Figure 1.**
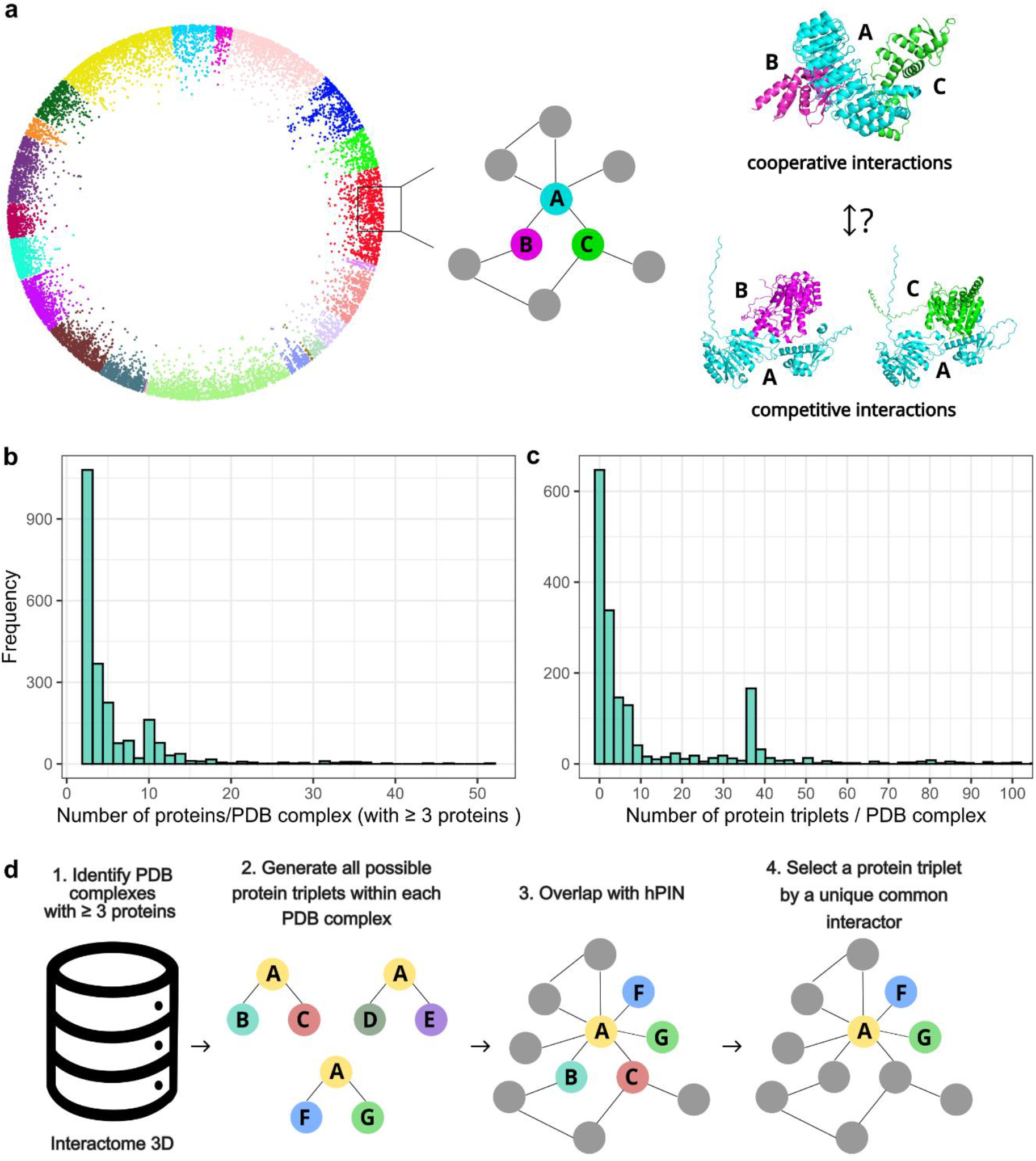
Construction of the human protein interaction network and identification of structurally supported cooperative triplets. **(a)** Overview of the high-confidence human protein–protein interaction network (hPIN) constructed from the HIPPIE database (confidence score ≥ 0.71) and embedded into two-dimensional hyperbolic space using the LaBNE+HM algorithm. Node colors represent functional clusters identified using the angular gap-based clustering method described in our previous study [18], where proteins grouped by angular proximity reflect functional similarity. **(b)** Distribution of the number of proteins per PDB complex with at least three unique proteins, based on residue-level annotations from Interactome3D. **(c)** Distribution of the number of structurally supported open triangle motifs (i.e., cooperative triplets) per PDB complex. A subset of complexes contributes disproportionately high numbers of triplets, reflecting structurally rich assemblies. **(d)** Mapping of structurally supported triplets into the hPIN. Only one cooperative triplet was retained per common interactor to ensure non-redundancy, resulting in a curated set of 211 cooperative triplets used for model training.

To investigate the diversity of protein structures involved in cooperative triplets, we analyzed residue-level annotations from the Interactome3D database [32]. We identified triplets within experimentally resolved complexes, focusing on open triangle configurations in which a central “common” protein binds two partners (V1 and V2) that do not interact directly. Such motifs are indicative of potential cooperative relationships, especially when the binding regions of the common protein are distinct or complementary. Across all PDB complexes containing at least three proteins, we observed a broad distribution in the number of proteins per complex, with most structures comprising fewer than 15 unique proteins (Figure 1b) [52].

These complexes were then computationally evaluated to identify open triangles. We found that while many PDB entries contained only a few open triangles, some complexes presented high structural complexity, producing up to 100 triplets (Figure 1c). Large well-studied assemblies such as ribosomes, proteasomes and chaperones, where many subunits are resolved together within the same structural experiment, result in large amounts of triplets [53,54]. Subunits of such large complexes participate in multiple triplets, which could be a source of bias in analyses of their properties.

After characterizing the structural diversity of complexes, we proceeded to map the identified triplets into the hPIN. This allowed us to extract structurally supported triplets with topological relevance, which served as cooperative examples for modeling. Specifically, we overlapped the common–V1 and common–V2 interaction pairs from each structurally validated triplet with the interactions present in the hPIN (Figure 1d). To avoid the redundancy that could be introduced by counting proteins from large complexes multiple times, as mentioned above, we retained only one cooperative triplet per common interactor. This filtering resulted in a final, non-redundant set of 211 cooperative triplets, distributed across 352 PDB complexes, each annotated with residue-level interface information derived from Interactome3D. These well-supported triplets formed the positive class used to train our classification model (Supplementary Table S4).

### Predicting cooperative triplets in the human protein interaction network

To systematically predict cooperative interactions among protein triplets, we constructed a dataset composed of positive and negative cases. The 211 structurally supported triplets identified from Interactome3D [32] served as the positive class, representing cooperative configurations in which a central protein (common interactor) binds two partners without a direct interaction between them (V1 and V2). As a “noisy” negative class, we selected open triangles from the hPIN that lacked structural support. These contain both positives and negatives. To eliminate any potential positional bias (for example, due to labels), we randomized the assignment of V1 and V2 within each triplet.

To build our model, we extracted for each triplet a set of topological and geometric features. Topological and hyperbolic features have been previously shown to be predictive of protein interaction with function in protein (de-)phosphorylation [16]. These included hyperbolic coordinates and centrality measures (degree, closeness, betweenness, eigenvector) for each of the three proteins in triplet, and distances, angular and radial differences for each pairwise relationship (common-V1, common-V2, V1-V2).

We also considered biological features: presence of a disordered region, and subcellular location, for each of the three proteins in triplet. From a biological perspective, we assumed that the formation of stable cooperative complexes requires that interacting proteins are co- localized in the same subcellular compartment [55]. In addition, proteins with well-defined structural domains are often capable of forming stable binding interfaces, making structural order a key indicator for such interactions [56].

In contrast, intrinsically disordered proteins (IDPs), or proteins with intrinsically disordered regions (IDRs), lack a well-defined tertiary structure under physiological conditions. While this structural flexibility can facilitate transient interactions such as in cellular signaling, it is generally less favorable for stable, multimeric complex formation [57]. The final feature matrix comprised 42 features per triplet (see Methods).

We trained multiple machine learning models - Random Forest (RF), Support Vector Machine (SVM), Logistic Regression, Decision Trees, and k-Nearest Neighbors (kNN) - and evaluated their performance using a 70/30 train-test split. Specifically, 70% of the dataset was used for model training and internal validation, while the remaining 30% was reserved for testing. To address the class imbalance between cooperative (positive) and competitive (negative) triplets, we applied random undersampling to the majority class in the training set prior to model training, resulting in a balanced dataset of 296 samples. For the training procedure, we performed 5-fold cross-validation repeated 10 times.

Among all tested classifiers, the Random Forest (RF) model (trained with 500 trees) achieved the highest performance, with a mean accuracy of 0.80, F1-score of 0.89, sensitivity of 0.80, and specificity of 0.80 on the held-out test set (Figure 2a). ROC curve analysis with an AUC of 0.88 (Figure 2b) indicated a superior ability to distinguish cooperative and competitive interactions between triplets of protein complexes. As our model predicts the interaction type within protein triplets, each unique triplet was evaluated once, resulting in a total of n = 17,165,561 triplet evaluations across the human protein interaction network. The model produces a probability score indicating whether a given triplet interaction is predicted to be cooperative or competitive. A total of 13,788,795 triplets received a score ≥ 0.5, and 5,769,218 triplets were classified with high confidence, receiving a score ≥ 0.9 (Supplementary Table S5).

**Figure 2.**
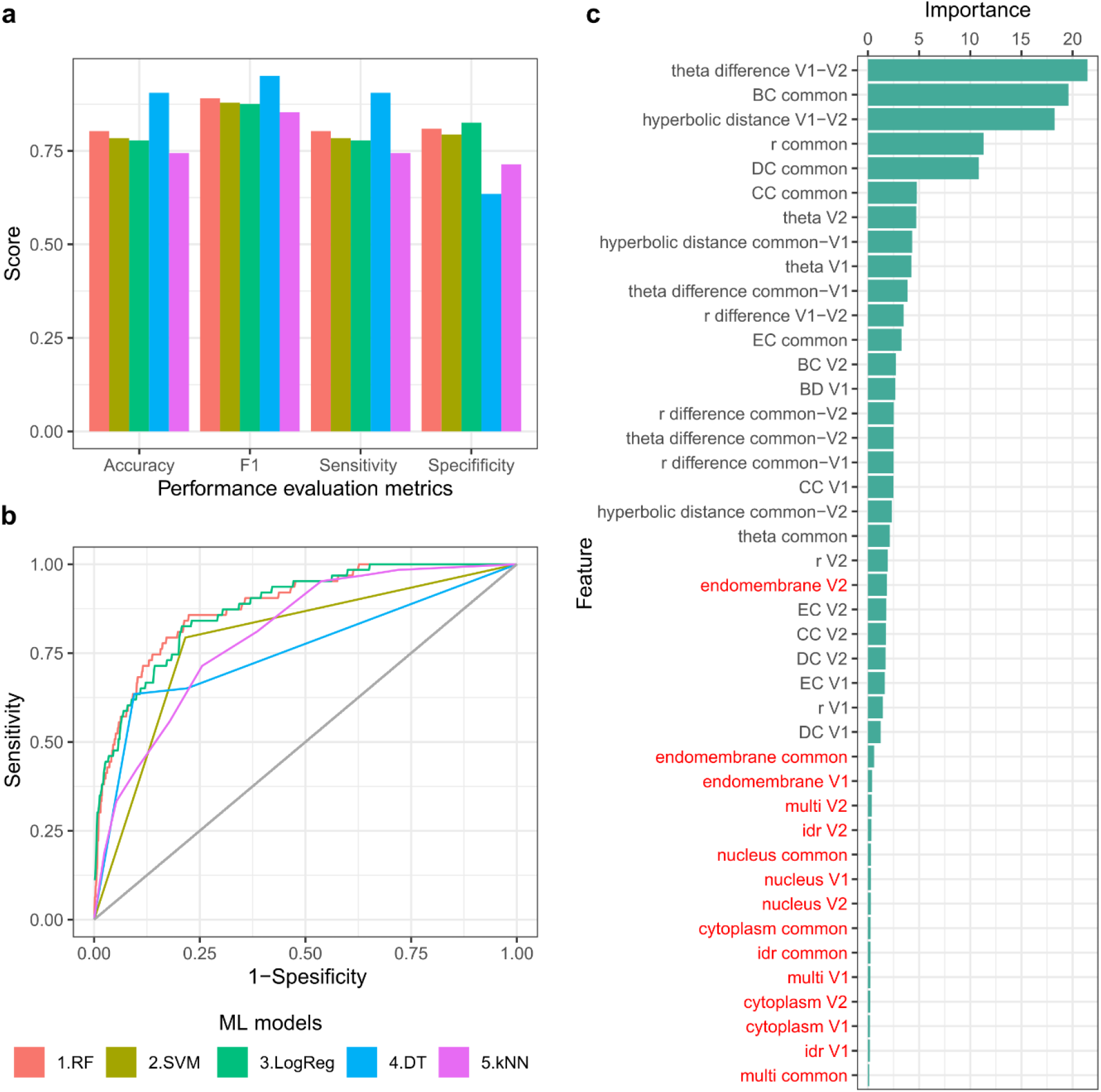
Predicting cooperative protein triplets in the human protein interaction network using machine learning. **(a)** Performance metrics (accuracy, F1-score, sensitivity, and specificity) of five machine learning models trained to classify cooperative versus competitive triplets. Random Forest (RF), Support Vector Machine (SVM), Logistic Regression (LogReg), Decision Trees (DT), and k-Nearest Neighbors (kNN). The Random Forest (RF) classifier showed the highest overall performance. **(b)** Receiver Operating Characteristic (ROC) curves for all models. The RF model achieved the best performance with an area under the curve (AUC) of 0.89, indicating strong discriminatory power. **(c)** Feature importance scores from the RF classifier, ranked by mean decrease in accuracy. The most informative features included the angular difference between V1 and V2, the betweenness centrality and hyperbolic distance of the common protein, and other geometric and topological features derived from the hyperbolic embedding and network structure. Biological features (labels marked in red) were less important.

To gain insight into which features were most predictive of cooperative interactions, we examined feature importance scores derived from the Random Forest model (Figure 2c). The most important feature was the angular difference between V1 and V2, closely followed by the hyperbolic distance between V1 and V2 proteins. The high importance of the angular and hyperbolic geometrical features aligns with previous observations that angular positioning in hyperbolic space captures the functional organization of the proteome [22].

Other top-ranking features included the betweenness (BC), degree (DC) and closeness (CC) centrality of the common protein as well as its radial coordinate. BC represents proteins that act like bridges in the network, DC values indicate how well a node is directly connected to the other nodes and CC measures the central position of the proteins in the network. Regarding the radial coordinate, in the hyperbolic embedding, proteins closer to the center (with lower radial values) are usually more connected in the network and often older in evolutionary terms.; they can be informative for predicting cooperation or competition between the proteins.

Together, these results highlight the value of hyperbolic embedding, which captures latent geometric and topological properties of the protein interaction network that cannot be easily inferred from biological features alone. Although biological attributes like subcellular localization and structural order add meaningful information, it was the geometric features derived from the network, especially angular and hyperbolic distances, that played the most critical role in distinguishing cooperative from competitive triplets. This demonstrates that embedding the hPIN in hyperbolic space enhances our ability to capture functional relationships and improve predictions of protein interaction types.

### Biological interpretation of predicted triplets

To investigate the biological relevance of our model’s predictions, we focused on the subset of predicted cooperative and competitive triplets where all three proteins had low degree centrality (degree ≤ 10).

This filtering step reduced the influence of highly connected hub proteins and allowed us to focus on triplets where cooperative assembly might be more functionally focused. From this subset, we retained 1,595 triplets and divided them into four quantiles based on their predicted probability scores, with Q1 representing low-scoring (competitive-like) and Q4 high-scoring (cooperative-like) predictions (Supplementary Table S6).

We first examined the sequence length of the proteins in each triplet under the assumption that longer proteins, particularly the common interactor, may contain more structural domains, enabling them to mediate cooperative interactions. Indeed, we observed a clear trend where the common protein’s sequence length increased across the quantiles (Figure 3a), suggesting that longer, possibly multi-domain proteins are more likely to participate in cooperative interactions. In contrast, the average length of V1 and V2 remained relatively constant, indicating that length-associated cooperative capacity is primarily associated with the common interactor.

**Figure 3.**
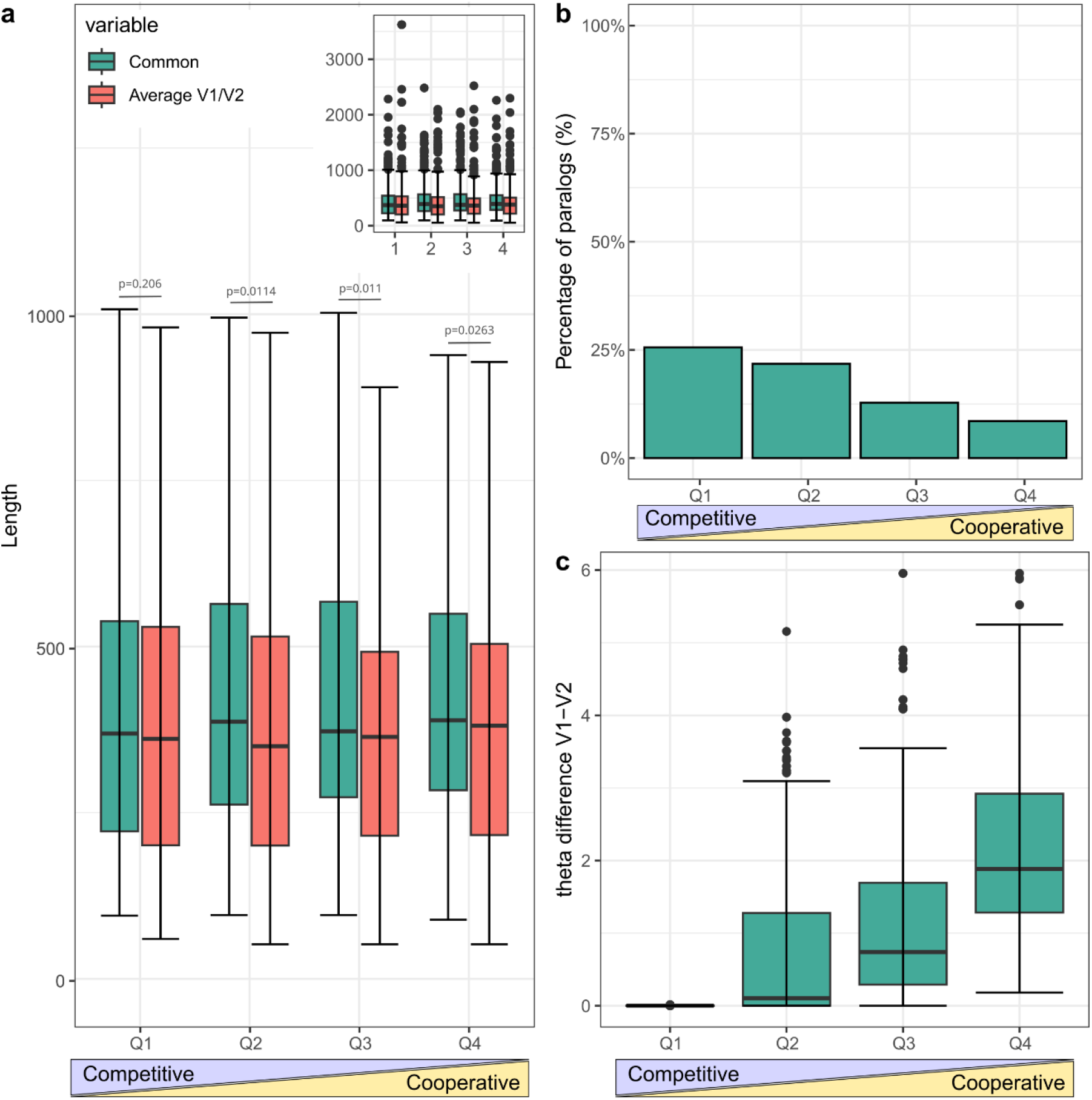
Biological interpretation of predicted triplets based on sequence length, paralogy, and angular separation. **(a)** Protein length distribution across predicted triplets, grouped into four quantiles based on model prediction scores (Q1 = low/competitive, Q4 = high/cooperative). The inset shows the outliers. The common protein in high-scoring (cooperative-like) triplets tends to be longer, suggesting a potential for multiple binding domains. **(b)** Proportion of triplets in which V1 and V2 are paralogs, across prediction quantiles. Paralogous pairs are more prevalent in low-scoring (competitive) triplets, supporting the idea that structurally similar proteins may compete for the same binding interface. **(c)** Distribution of angular (theta) difference between V1 and V2 across the four quantiles. Smaller angular differences are associated with low-scoring triplets, indicating that functionally similar partners tend to compete, whereas cooperative triplets involve functionally diverse proteins.

Next, we assessed the paralogous relationship between V1 and V2 in each triplet. The hypothesis was that paralogous proteins may compete for the same binding site on the common interactor due to structural and functional similarity. Consistent with this, we found that the proportion of paralogous V1–V2 pairs was highest in Q1 and decreased steadily toward Q4 (Figure 3b), supporting the idea that paralog pairs are more likely to be involved in competitive interactions.

Finally, we examined the angular (theta) difference between V1 and V2, the top-ranked feature from our model. Triplets in Q1 exhibited the lowest angular separation, while those in Q4 had significantly higher theta differences (Figure 3c). This aligns with the analysis that cooperative interactions are more likely to involve functionally diverse proteins, reflected in a broader separation in angular space. In contrast, small angular differences (indicating functional similarity) were associated with predicted competitive interactions, reinforcing the idea that competition is more likely when both proteins occupy similar regions in the hyperbolic space.

### Structural evaluation of predictions using AlphaFold

To evaluate whether our predictions are meaningful from a structural point of view, we selected four representative triplets for structural modeling: two from the top-ranked cooperative predictions and two from the bottom-ranked competitive predictions, based on their respective classification scores. For each triplet, we generated AlphaFold 3 models of the two binary interactions (common–V1 and common–V2) yielding eight predicted complexes in total (Figure 4).

**Figure 4.**
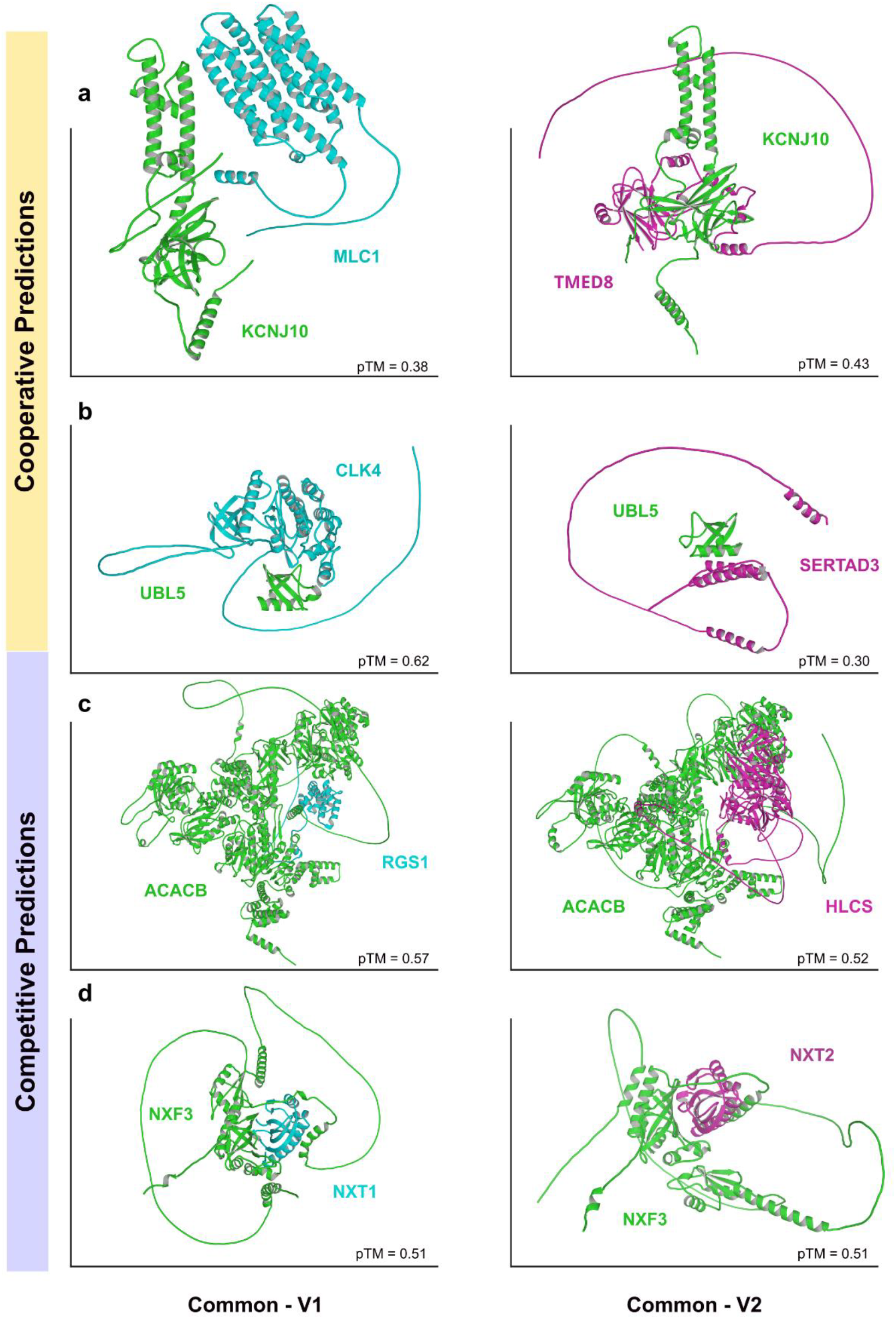
Structural validation of model predictions using AlphaFold. Representative examples of predicted cooperative (top two rows) and competitive (bottom two rows) triplets. Each panel shows a binary complex between the common protein and one of its partners, modeled with AlphaFold 3. In cooperative cases, the two partners bind to distinct, non-overlapping regions on the common protein: **(a)** Q6PL24 (TMED8) and Q15049 (MLC1) binding to MLC1P78508 (KCNJ10); **(b)** Q9HAZ1 (CLK4) and Q9UJW9 (SERTAD3) binding to Q9BZL1 (UBL5). In competitive cases, both partners share overlapping or adjacent binding interfaces with the common protein: **(c)** Q08116 (RGS1) and P50747 (HLCS) binding to O00763 (ACACB); **(d)** Q9UKK6 (NXT1) and Q9NPJ8 (NXT2) binding to Q9H4D5 (NXF3).

We report pTM values as a general measure of model confidence, even though they focus on overall structure instead of the interaction interface. Therefore, to assess the nature of the interfaces, we manually reviewed the predicted structures using PyMOL, analyzing the positioning of residues at the binding interface. Triplets with non-overlapping binding sites on the common protein were interpreted as cooperative, while overlapping or adjacent binding surfaces indicated potential competition between V1 and V2.

The triplet composed of P78508 (*KCNJ10*) binding to Q6PL24 (*TMED8*) and Q15049 (*MLC1*) was predicted by our model to be cooperative, based on a high classification score and structural evidence (Figure 4a). Although AlphaFold has generally problems folding transmembrane proteins because it does not consider the membrane, we can see that transmembrane protein KCNJ10 can interact with MCL1, which is another transmembrane protein, using contacts between transmembrane helices, whereas its interaction with TMED8 happens via its cytoplasmic domain, allowing a cooperative interaction with these two proteins. *KCNJ10* encodes Kir4.1, a potassium channel crucial for ion homeostasis in glial cells, with mutations linked to EAST syndrome [58]. *MLC1* is expressed in astrocytes and contributes to water and ion regulation at the blood-brain barrier; its dysfunction causes megalencephalic leukoencephalopathy [59]. While *TMED8* (Transmembrane p24 Trafficking Protein Family Member 8) is named as part of the TMED family, its direct involvement in protein trafficking is not well-established in scientific literature. A review by Zhang et al. discusses the TMED family’s role in intracellular protein transport, particularly between the endoplasmic reticulum (ER) and Golgi apparatus. However, the authors note that *TMED8* differs structurally from other TMED proteins: it lacks a transmembrane domain and two conserved cysteine residues, and therefore may not function as a typical p24 family member involved in vesicular trafficking [60]. Despite lacking a transmembrane domain and conserved cysteine residues typical of TMED family proteins, the presence of *TMED8* within the same triplet as *KCNJ10* and *MLC1* (two astrocyte-associated proteins involved in ion regulation) raises the possibility that *TMED8* may contribute to cooperative function through an alternative or indirect role in membrane-associated processes.

A similar pattern was observed in the second cooperative example of Q9BZL1 (*UBL5*) binding to Q9HAZ1 (*CLK4*) and Q9UJW9 (*SERTAD3*), where the two partners were positioned on opposing sides of the common interactor, with no interface overlap observed in the AlphaFold models. UBL5 is a ubiquitin-like protein involved in mRNA splicing and sister chromatid cohesion, functioning through interactions with spliceosomal proteins and contributing to post-transcriptional gene regulation [61]. While UBL5 shuttles between nucleus and cytoplasm, CLK4 and SERTAD3 are both nuclear proteins. CLK4 is a dual-specificity protein kinase that phosphorylates serine/arginine-rich (SR) proteins, which are core components of the spliceosome. Through this activity, CLK4 modulates alternative splicing and contributes to transcriptomic diversity [62]. The third protein in this triplet, SERTAD3, is a transcriptional co-regulator implicated in cell proliferation, apoptosis, and responses to interferon signaling. It has been shown to influence innate immune responses and may act downstream of p53- related pathways in cancer and antiviral defense [63]. Together, these proteins are associated with different aspects of gene expression regulation, supporting the possibility of cooperative interaction. The link between UBL5 and CLK4 in RNA processing may work together with SERTAD3’s role in transcriptional regulation, suggesting that this triplet could function as part of a system that coordinates gene expression control (Figure 4b).

In contrast, two additional triplets classified by the model as competitive provide further structural and biological support for mutually exclusive interactions. The first competitive triplet consists of O00763 *(ACACB*) interacting with Q08116 (*RGS1*) and P50747 (*HLCS*). ACACB encodes acetyl-CoA carboxylase beta, a key mitochondrial enzyme in fatty acid metabolism that catalyzes the conversion of acetyl-CoA to malonyl-CoA [64]. *RGS1* is selectively expressed in lymphoid tissues and functions as a GTPase-activating protein (GAP), thereby attenuating G-protein–mediated signaling in immune cells [65]. RGS1 is not a mitochondrial protein and the only evidence of its interaction with mitochondrial ACACB is from two large high-throughput studies [66,67]. The other interaction is well established: holocarboxylase synthetase (*HLCS*) is an essential enzyme that catalyzes the biotinylation of multiple carboxylases involved in fatty acid synthesis, amino acid metabolism, and gluconeogenesis, particularly of ACACB [66]. Structural modeling of this triplet revealed partial overlap between the binding sites of RGS1 and HLCS on ACACB (Figure 4c). Combined with their distinct biological functions (signal transduction for RGS1 and metabolic regulation for HLCS), this observation would support the prediction that a putative interaction of RGS1 with ACACB would compete for binding with HLCS, rather than forming a cooperative complex.

In the second competitive triplet, Q9H4D5 (*NXF3*) binds to Q9UKK6 (*NXT1*) and Q9NPJ8 (*NXT2*). These interactions are well supported by experimental evidence [68]. The three proteins are associated with nuclear mRNA export. NXF3 is a member of the nuclear RNA export factor family, mediating mRNA export from the nucleus to the cytoplasm [69]. NXT1 and NXT2 are closely related paralogs involved in RNA export, functioning as cofactors in both RAN- and CRM1-dependent nuclear export pathways [70]. Notably, because NXT1 and NXT2 are paralogs, they share high structural similarity, which likely enables both proteins to bind the same domain on NXF3. Structural modeling confirmed that NXT1 and NXT2 bind to nearly identical interface regions in the NTF2 domain of NXF3, suggesting strong competition for binding (Figure 4d). A similar interface of interaction can be found in an experimentally-solved structure between NXT1 and the NTF2 domain of NXF1 (a paralog of NXF3) (PDB: 6E5U;[71]).

While our model generally performs well in distinguishing cooperative from competitive triplets, there are cases where caution is warranted. For example, the triplet with Q7Z6M4 (*MTERF4*) binding to P51460 (*INSL3*) and Q9BWT6 (*MND1*) was predicted as cooperative based on the classification score. However, AlphaFold structural modeling suggests that INSL3 and MND1 bind to overlapping regions on the surface of MTERF4, suggesting potential competition for the same interface rather than cooperative engagement (model not shown). This discrepancy suggests that relying only on network topology and geometry can limit the model’s accuracy in certain cases. However, it also highlights the value of using structural modeling afterward to validate predictions and gain deeper insight into cases that are less clearly defined.

## Conclusions

In this study, we systematically explored cooperative and competitive interactions among protein triplets by combining structural data, network topology, hyperbolic embeddings, and machine learning-based prediction. Our results provide novel insights into the structural and functional determinants that differentiate cooperative from competitive assemblies in the human protein interaction network (hPIN). By embedding the hPIN into hyperbolic space, we were able to capture latent geometric properties of the network that traditional biological or topological features alone could not fully reveal. The angular separation between proteins emerged as the most informative feature for predicting interaction type, highlighting the value of hyperbolic geometry in modeling functional organization within complex biological networks. This finding is consistent with previous work showing that angular proximity in hyperbolic space reflects functional similarity, while radial positioning captures evolutionary and topological hierarchy.

Our machine learning models, particularly the Random Forest classifier, demonstrated strong performance in distinguishing cooperative from competitive triplets, achieving high accuracy, F1-score, and AUC values. We observed that biological context such as subcellular localization and the presence of intrinsically disordered regions did not contribute to predictive success as much as geometrical and topological features. However, biologically meaningful patterns were recovered regarding biological properties not used for training such as protein length and paralogy: cooperative triplets tended to involve longer common proteins, reflecting the presence of multiple or independent binding sites, while competitive triplets often involved paralogous proteins competing for the same interaction surface.

Validation through AlphaFold 3 demonstrated that cooperative interactions involve distinct interfaces, while competitive interactions are characterized by significant interface overlap. The identification of cooperative modules involved in processes such as ion transport, vesicle trafficking, and gene expression strengthens the biological significance of our results. Conversely, competitive triplets enriched for paralogous proteins and redundant pathways showcase the complexity of competition-driven regulatory mechanisms in cellular signaling and molecular transport.

In conclusion, our study demonstrates that integrating network-based features, hyperbolic embeddings, and machine learning provides a powerful framework for investigating higher-order protein interactions. Our findings demonstrate the potential of geometric approaches as powerful tools for exploring the complexity of biological networks.

## Data availability

All supplementary data supporting the findings of this study, including node- and edge-level features, structural annotations, and prediction scores, are available at: https://github.com/avagiona/Cooperative-Competitive-interactions-within-protein-triplets

## SUPPLEMENTARY TABLES

**Supplementary Table S1**. Nodes of the hPIN. Columns indicate protein identifiers (UniProtKB), hyperbolic coordinates (r, theta), centrality measures (Degree Centrality—DC, Closeness Centrality—CC, Betweenness Centrality—BC, and Eigenvector Centrality—EC), intrinsic disorder region presence (idr; yes=1, no=0) and subcellular localization (nucleus, cytoplasm, endomembrane, multi-localized proteins).

**Supplementary Table S2**. Edges of the hPIN. Columns indicate protein identifiers (UniProtKB; V1, V2), hyperbolic distance (hd), r difference (rd) and angular difference (thetad) derived from the hyperbolic embedding.

**Supplementary Table S3**. Structurally supported protein triplets annotated using Interactome3D. Columns indicate the triplet identifier (Triplet_ID), associated PDB structure (PDB_ID), the common protein and its interactors (Common, V1, V2), and the regions of interaction between the common protein and each interactor (region_common_V1, region_common_V2), specifying the PDB chain and residues involved.

**Supplementary Table S4**. Feature matrix used for the classification of cooperative versus non-cooperative protein triplets. Each row represents a protein triplet, including the triplet type (cooperative or not), the protein identifiers (common, V1, V2), and hyperbolic embedding-based features (r, theta, hyperbolic distance—hd, radial distance—rd, angular difference— theta_d) for all proteins and their pairwise combinations. Node-level features (degree, closeness, betweenness, eigenvector centralities), intrinsic disorder region information (idr), and subcellular localization (nucleus, cytoplasm, endomembrane, multi-localized) are provided for each protein individually (common, V1, and V2).

**Supplementary Table S5**. Predictions of cooperative and non-cooperative protein triplets with associated scores. Columns indicate whether structural evidence supports the interaction (cooperative: 1 for structurally validated, 0 otherwise), the protein identifiers (common, V1, V2), and the classification score assigned by the predictive model.

**Supplementary Table S6**. Predictions of cooperative and competitive protein triplets based on low-degree proteins (DC≤10) triplets with structural validation and feature annotations. Columns include the validation label based on structural evidence (cooperative: 1 for structurally validated, 0 otherwise), protein identifiers (common, V1, V2), angular difference between V1 and V2 in hyperbolic space (thetad_V1_V2), degree centralities of each protein (degree_common, degree_V1, degree_V2), prediction score (score), quantile (quantile), presence of paralogous relationships (paralog), sequence lengths of each protein (common_length, V1_length, V2_length), and the average length of V1 and V2 (average_V1_V2).

